# *Uvariopsis ebo* (Annonaceae) a new, Critically Endangered tree species from the Ebo Forest, Cameroon and a key to the Cameroonian species of *Uvariopsis*

**DOI:** 10.1101/2021.03.26.437154

**Authors:** George Gosline, Martin Cheek, Jean Michel Onana, Eric Ngansop, Xander van der Burgt, Leo-Paul Dagallier

## Abstract

A new species to science of evergreen forest tree, *Uvariopsis ebo* (Annonaceae) is described, illustrated, mapped, and compared morphologically with the other cauliflorous species of the genus. Restricted so far to a single site in evergreen lowland forest in the Ebo Forest, Yabassi, Littoral Region, Cameroon, this species is Critically Endangered using the IUCN 2012 standard, because the forest habitat of this species remains unprotected, and there exists threats of logging and conversion to plantations. This species adds to the growing list of threatened species resulting from anthropogenic pressure on Cameroon forests. Observations on the unusual corolla structure of the new species are made. A revised key to the 14 Cameroonian species of *Uvariopsis* is presented. Notes are given on other narrowly endemic and threatened species in the Ebo forest area, a highly threatened centre of diversity in Littoral Region, globally important for conservation.

## Introduction

The new species reported in this paper was discovered as a result of a long-running survey of plants in Cameroon to support improved conservation management. The survey is led by botanists from the Royal Botanic Gardens, Kew and IRAD (Institute of Agricultural Research for Development)-National Herbarium of Cameroon, Yaoundé. The study has focussed on the Cross-Sanaga interval (Cheek *et al*. 2001, 2006) which contains the area with the highest plant species diversity per degree square in tropical Africa (Barthlott *et al*. 1996). The herbarium specimens collected in these surveys formed the foundations for a series of Conservation Checklists (see below). So far, over 100 new species and several new genera have been discovered and published as a result of these surveys, new protected areas have been recognised and the results of analysis are feeding into the Cameroon Important Plant Area programme (https://www.kew.org/science/our-science/projects/tropical-important-plant-areas-cameroon), based on the categories and criteria of Darbyshire *et al*. (2017).

In connection with preparation of a Conservation Checklist of the plants of the Ebo Forest, Littoral Region, Lorna MacKinnon, a volunteer botanical assistant, made a plant collecting expedition to the Ebo Forest in 2008. Among the collected plant specimens and photographs that she made, we identified an *Uvariopsis* Engl. & Diels (*MacKinnon* 51, Fig. 1 & 2) which resembled no other species known in the genus. In this paper we test the hypothesis that it is a new species to science, and name the new species as *Uvariopsis ebo*.

**Fig. 1.**
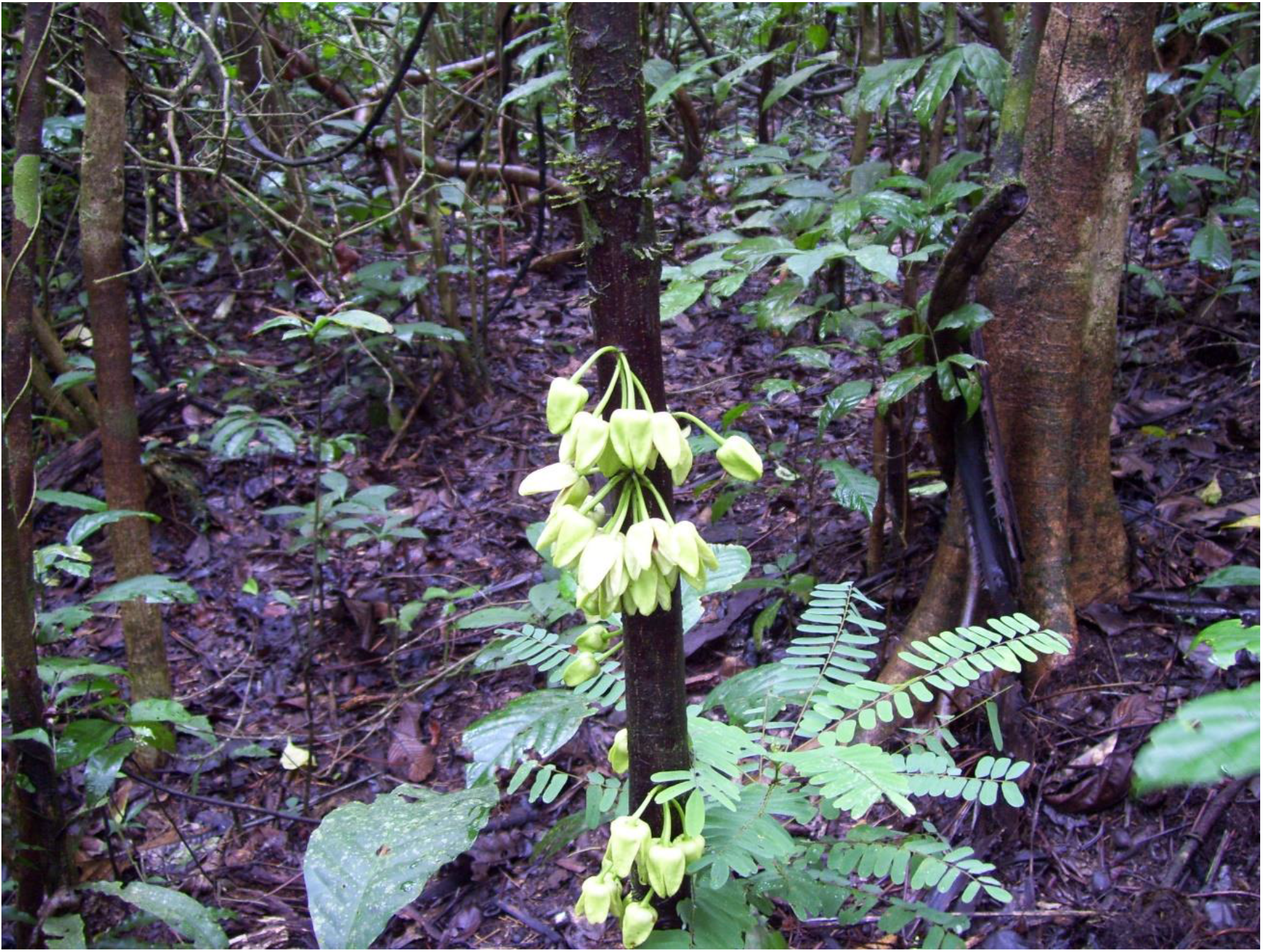
*Uvariopsis ebo*. Cauliflorous inflorescences. *MacKinnon* 51(K). Photo Lorna MacKinnon.

**Fig. 2.**
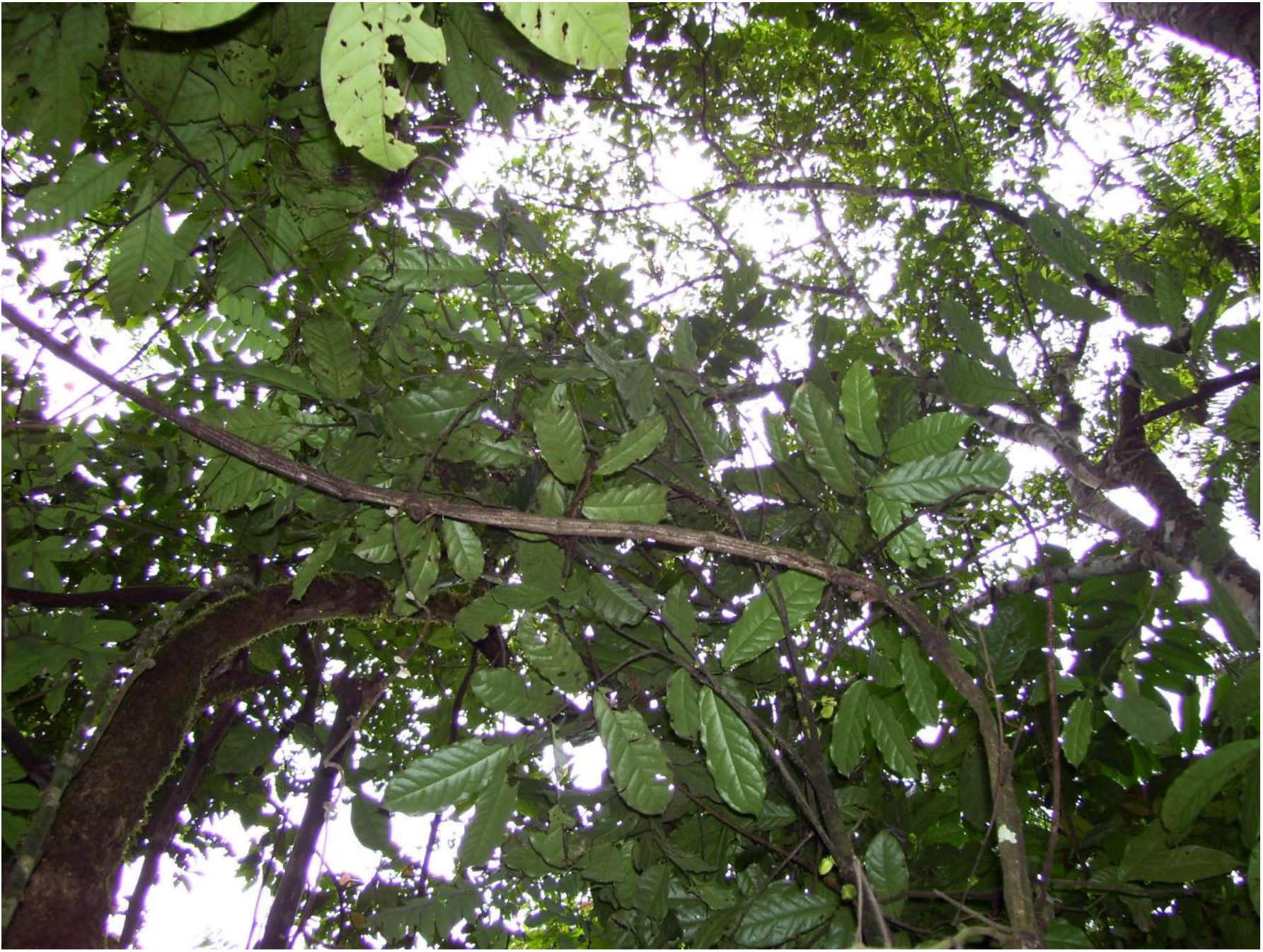
*Uvariopsis ebo*. Trunk apex with cauliflorous flowers and canopy. *MacKinnon* 51(K). Photo Lorna MacKinnon.

*Uvariopsis* (Annonaceae) is a highly distinctive and easily recognised genus, since most of its species have unisexual flowers, a calyx with two basally connate sepals, and the petals in a single whorl of four (very rarely three, see below). Annonaceae are otherwise characterised by bisexual flowers with trimerous perianths. Most species of the genus are cauliflorous small trees, the flowers being produced from the trunk, although some species are ramiflorous or bear axillary flowers (Kenfack *et al*. 2003, Couvreur *et al*. 2021).

Nineteen species are currently accepted in *Uvariopsis*. Five species have been published in the 21^st^ Century: *Uvariopsis korupensis* Gereau & Kenfack (Gereau & Kenfack 2000), *U. submontana* Kenfack, Gosline & Gereau (Kenfack *et al*. 2003) and *U. etugiana* Dagallier & Couvreur (Couvreur *et al*. 2021) all from Cameroon, *U. citrata* Couvreur & Niang. (Couvreur & Niangadouma 2016) from Gabon and *U. lovettiana* Couvreur & Q. Luke (Couvreur & Luke 2010) from Tanzania. In addition a sixth species, *Uvariopsis tripetala* (Baker f.) G.E.Schatz, was transferred to the genus from the monotypic *Dennettia* Baker f. (Kenfack *et al*. 2003). The genus is centred in Cameroon, where 13 of the 19 species occur, followed in species diversity by Gabon, with six species. The most widespread species of the genus is *Uvariopsis congensis* Robyns & Ghesq. Which occurs from Cameroon to S.Sudan, Zambia and Kenya. Several species are rare, being known from only one or two specimens and have restricted ranges, these include *U. etugiana* (Cameroon endemic) and *U. citrata* (Cameroon & Gabon), both known from two specimens, and *U. sessiliflora* (Mildbr. & Diels) Robyns & Ghesq. endemic to Cameroon and known from a single specimen.

The genus is distributed throughout continental tropical African evergreen forests, from Guinea in the West to Tanzania in the east, and as far south as northern Zambia. The species usually occur at low altitude, exceptions being *U. submontana* and *U. lovettiana* which occur in submontane or cloud forest in Cameroon and Tanzania respectively (Kenfack *et al*. 2003; Couvreur & Luke 2010). Species are usually small trees in high quality, undisturbed forest and appear not to be pioneers. They usually occur at low frequency. For example, in the Mefou Proposed National Park of Central Region Cameroon, only a single mature individual with one juvenile of one species of the genus, *U. solheidii* (De Wild.) Robyns & Ghesq., was found in the course of many weeks of botanical surveys by numerous botanists collecting thousands of specimens (Cheek *et al*. 2011: 123). However, in some rare ecological circumstances, some species can become locally dominant e.g., *U. tripetala* in the understorey of maritime lowland evergreen inselberg forest in Guinea (Couch *et al*. 2019: 41), and also *U. congensis* locally subdominant in forests in western Uganda where it flowers synchronously and is dispersed by primates (Dominy & Duncan 2005; Gosline pers. obs. 2016).

## Materials and Methods

Herbarium citations follow Index Herbariorum (Thiers *et al*. 2020). Specimens were viewed at EA, K, P, WAG, and YA. The National Herbarium of Cameroon, YA, was searched for additional material of the new species, but without success. Images for specimens at WAG were studied at https://bioportal.naturalis.nl/?language=en and those from P at https://science.mnhn.fr/institution/mnhn/collection/p/item/search/form?lang=en_US. We also searched JStor Global Plants (https://plants.jstor.org/ accessed March 2021) for additional material, and finally the Global Biodiversity Facility (GBIF, www.gbif.org accessed March 2021). Binomial authorities follow the International Plant Names Index (IPNI, 2020). The conservation assessment was made using the categories and criteria of IUCN (2012). Herbarium material was examined with a Leica Wild M8 dissecting binocular microscope fitted with an eyepiece graticule measuring in units of 0.025 mm at maximum magnification. The drawing was made with the same equipment using Leica 308700 camera lucida attachment. The botanical terms follow Beentje & Cheek (2003), and format of the description follow the conventions of Kenfack *et al*. (2003) and Couvreur & Luke (2010). The map was made using SimpleMappr (https://www.simplemappr.net).

## Results

Because it has leaves exceeding 15 cm long, cauliflorous flowers with pedicels exceeding 10 mm long, petals exceeding 7 mm long, *MacKinnon* 51, described below as *Uvariopsis ebo*, keys out in the key to the species of the genus in Couvreur *et al*. (2021) to couplet 12, leading to *U. dioica* (Diels) Robyns & Ghesq. (flower buds globose) and *U. solheidii* (flowers buds conical). Of these two choices, our material is closest to the second of these leads, the buds being narrowly ovoid. The two species are similar, but can be separated using the differential characters in Table 1, and the new species can be separated from all other cauliflorous species of *Uvariopsis* using the key (both presented below).

**Table 1.** Differential characters separating *Uvariopsis solheidii* and *U.ebo*. Data for *Uvariopsis solheidii* taken mainly from Couvreur *et al*. (2021) and Cheek *et al*. (2011).

| Character | *Uvariopsis solheidii* | *Uvariopsis ebo* |
| --- | --- | --- |
| Indumentum of stem, petiole and abaxial midrib | Tomentose | Glabrous |
| Leaf-blade dimensions | 16.6 – 29 x 5 – 9.5 cm | 17.7 – 20.3 (– 23) x (6.4 – ) 7 – 7.9 cm |
| Number of secondary nerves on each side of midrib | 8 – 13 | 5 – 8 |
| Number of flowers per inflorescence | 2 – 3 | 4 – 7 |
| Petal dimensions | 7 – 10 x 2.5 – 5 mm | (14 – ) 16 x (5.5 – ) 9 mm |
| Petal colour | Wine brown | Yellow-green |

### Key to the species of *Uvariopsis* in Cameroon

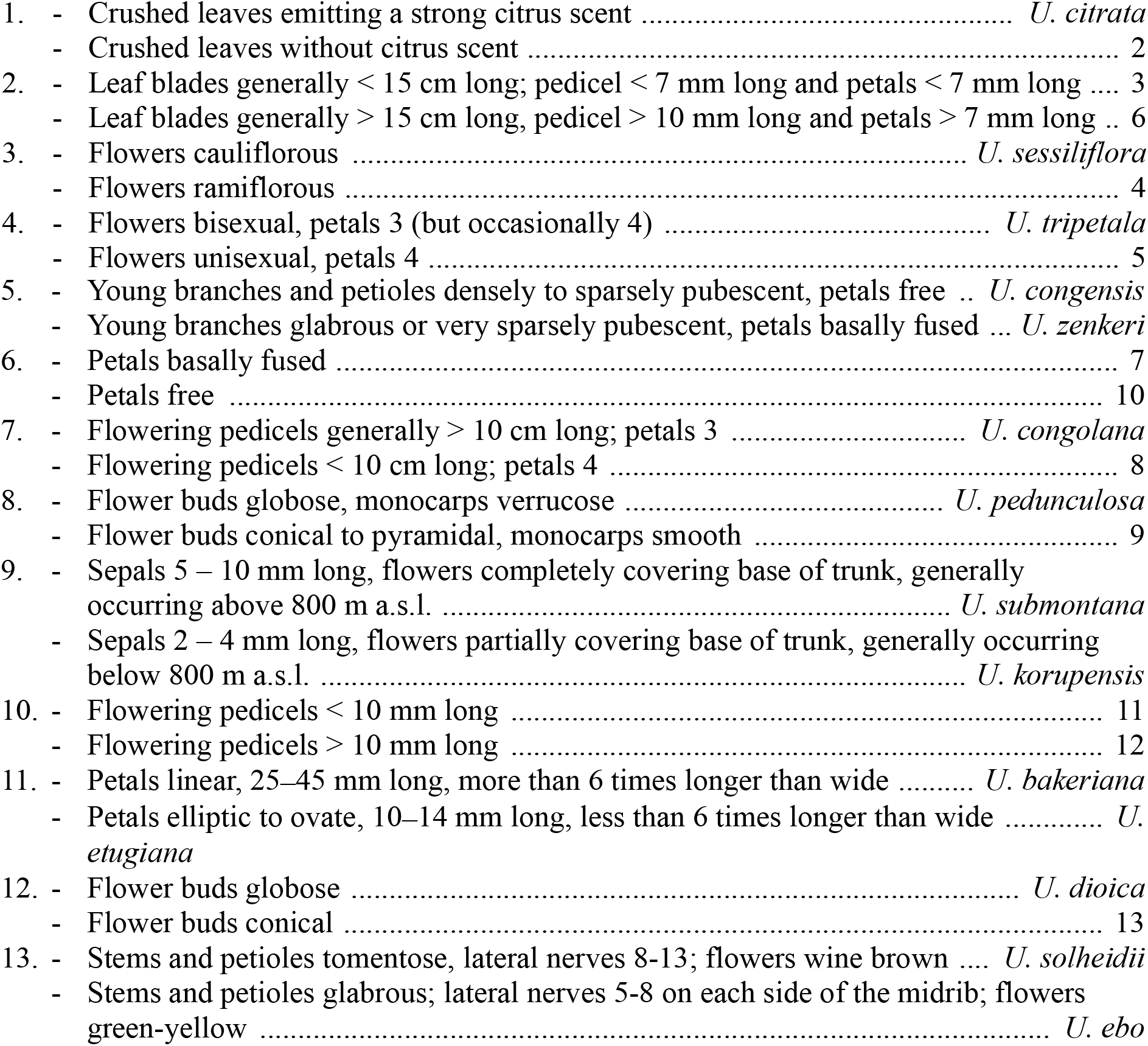

***Uvariopsis ebo** Cheek & Gosline* **sp. nov**. Type: Cameroon, Littoral Region, Ebo Forest, 4° 20’ 44” N, 10° 24’ 33”E, 849 m alt. Dicam Trail 2000 m from Bekob camp, male fl. 25 March 2008, *MacKinnon* 51 (holotype K001381842; isotypes YA, MO).

*Cauliflorous, probably monoecious understorey tree* 3 – 4 m tall. Trunk terete, lacking flutes or prop roots, 1.8 – 2.5 cm diameter at 1.5 m above the ground, bark smooth, dark-brown (Fig. 1), the crown sparsely branched (Fig. 2). Leafy stems with 3 – 4 leaves per season’s growth, terete, internodes (1.2 −) 1.5 – 2.8 (− 4.3) cm long, 0.15 – 0.2 cm diam., pale yellow-green, later orangish brown, glabrous. Axillary buds dome-shaped 0.5 – 0.75 x 1 mm, bud-scales numerous, linear, spreading, densely hairy, hairs simple, appressed hairy c. 0.5 mm long, colourless or red brown. *Leaves* distichous, held in the horizontal plane, lacking scent when crushed (collection data, *MacKinnon* 51), blades oblanceolate 17.7 – 20.2 (− 23) x (6.4 −) 7 – 7.9 cm, acumen narrowly triangular (0.5 −) 1 – 1.3 cm, base broadly acute with convex edges, minutely cordate, blade mounted above petiole, margins undulate-sinuous (live & dried), midrib impressed on adaxial surface, inconspicuous, below a grooves; on abaxial surface subcylindric, 1 – 1.2 mm diam., conspicuous; secondary veins 5 – 8 (− 9) on each side of the midrib, brochidodromous, arising at c. 50° from the midrib, initially straight, then curving in the outer third, uniting with the secondary nerve above to form a looping inframarginal nerve, attaining 3 – 4 mm from the margin; intersecondary nerves sometimes present, tertiary nerves raised, conspicuous, forming a reticulum with cells 4 – 5 mm long, quaternary nerves inconspicuous; glabrous (except in bud when densely orange-brown hairy, hairs 0.1 mm long). Petiole stout, shallowly canaliculate, 4 (− 5) mm long, 1.9 – 2.1 mm diam., narrowing at base and apex, the adaxial groove shallow, c. 0.5 mm wide, glabrous.

*Female inflorescences* unknown. *Male inflorescences* cauliflorous, scattered along the trunk from ground level to the top of the trunk 2.5 – 3 m above the ground, each 3 – 7-flowered (Fig. 1 & 2). Peduncles patent, c. 2 x 2 mm, pale brown, glabrous, bearing sub-umbellate, radiating, 1-flowered partial-peduncles. Partial-peduncles 0.5 – 2 x 0.9 – 1.2 mm, terminating in 1 – 2 bracts subtending a pedicel. Bracts oblong-elliptic, 1.5 x 0.5 – 0.6 mm, apex acute, outer surface about 50% covered in appressed white hairs c. 0.15 – 0.2 mm long, inner surface glabrous. Pedicels 1.8 – 2.5 cm long, 0.1 cm diam., articulated with the partial-peduncle, with (0 −) 1 (− 2) scattered, bracteoles in the proximal few mm. Bracteoles similar to the bracts, ovate-oblong, shortly sheathing, (1 −) 1.25 x 1 mm, outer surface with sparse scattered simple appressed translucent hairs 0.05 – 0.2 mm long. buds narrowly ovoid, 16 x 11 mm. *Sepals* 2, opposite, drying pale brown, reflexed, *Male flowers* semi-orbicular, 1 – 1.5 mm long, 2.1 – 2.5 mm wide, glabrous. *Petals* 4, uniseriate, free, yellow-green when live (Fig. 1), drying black, lanceolate-oblong, (14 −)16 x (5.5 −)9 mm, apex rounded, base rounded, outer surface sparsely and inconspicuously hairy,), hairs 7 – 9 per mm^2^ (5% of surface covered), simple, translucent, appressed, 0.1 mm long, apices rounded. Inner surface of petals with a shallow elliptic-oblong excavation c. 8 x 5 mm, the margin of the excavation raised, the apex with a ridge extending along the midline to the petal apex, glabrous apart from a few scattered erect, minute white hairs 0.05 mm long at the excavation apex. Staminal dome 3.5 – 4 mm long, 3.5 – 4 mm diam., consisting of stamens and a receptacular torus. Stamens shortly cylindrical-angular, 0.5 x 0.1 (− 0.2) mm, connective surrounded by four longitudinal anther cells, each exceeding the connective, in plan view the stamen is 4-lobed. Apical connective appendage absent. Female flowers, fruit and seed unknown. Fig. 1 & 2.

#### RECOGNITION

Similar to *Uvariopsis solheidii* (De Wild.) Robyns & Ghesq., differing in the stem, petioles and abaxial midrib glabrous (versus tomentose); number of secondary nerves on each side of the midrib 5 – 8 (versus 8 – 13); petals yellow-green, (14 −) 16 x (5.5 −) 9 mm (versus wine brown, 7 – 10 x 2.5 – 5 mm). Additional differential characters are given in Table 1, above.

#### DISTRIBUTION

Cameroon (Map1), endemic to the Ebo Forest of the Littoral Region on present evidence.

#### SPECIMENS EXAMINED

Cameroon, Littoral Region, Yabassi, Ebo Forest, 4° 20’ 44” N, 10° 24’ 33”E, 849 m alt. Dicam Trail 2000 m from Bekob camp, male fl. 25 March 2008, *MacKinnon* 51 (holotype K001381842; isotypes YA, MO).

#### HABITAT

*Uvariopsis ebo* is so far only known from lower submontane forest (849 m elev.). below the elevation for the upper montane forest indicator species *Podocarpus latifolius* (Thunb.) R.Br. ex Mirb. The geology is ancient, highly weathered basement complex, with some ferralitic areas in foothill areas which are inland, c. 100 km from the coast. Altitude varies from c. 200 m to 1200 m elevation. The wet season (successive months with cumulative rainfall >100 mm) falls between March and November and is colder than the dry season. Average annual rainfall at Bekob measured 2010 – 2016 is 2336 mm (Abwe, Ebo Forest Research Programme, Cameroon pers. comm., Abwe & Morgan 2008, Cheek *et al*. 2018a).

#### CONSERVATION STATUS

*Uvariopsis ebo* is currently known from a single specimen at a single location inside the mid-eastern part of the Ebo Forest. Less than 50 mature individuals have been observed (Bethan Morgan pers. comm. to Cheek, March 2021), despite the species being highly conspicuous in flower (Fig. 1) and situated on a major footpath close to a research camp used by many biologists over the last 15 years.

Since 2006, botanical surveys have been mounted almost annually, at different seasons, over many parts of the formerly proposed National Park of Ebo. About 2500 botanical herbarium specimens have been collected, but this species has been seen not yet been seen elsewhere in the c. 2000 km^2^ of the Ebo Forest. However, the area outside the two research camps, especially the western edge, has not been fully surveyed for plants. While it is likely that the species will be found at additional sites within the Ebo Forest, there is no doubt that it is genuinely range-restricted as are some other species of *Uvariopsis* in Cameroon (see introduction). Botanical surveys and other plant studies for conservation management in forest areas north, west and east of Ebo, and just over the border in Nigeria, resulting in tens of thousands of specimens being collected and identified have failed to find any additional specimens of this species (Cheek *et al*. 1996; Cable & Cheek 1998; Cheek *et al*. 2000; Maisels *et al*. 2000, Chapman & Chapman 2001, Harvey *et al*. 2004; Cheek *et al*. 2004; Cheek *et al*. 2010; Harvey *et al*. 2010; Cheek *et al*. 2011).

**Map 1.**
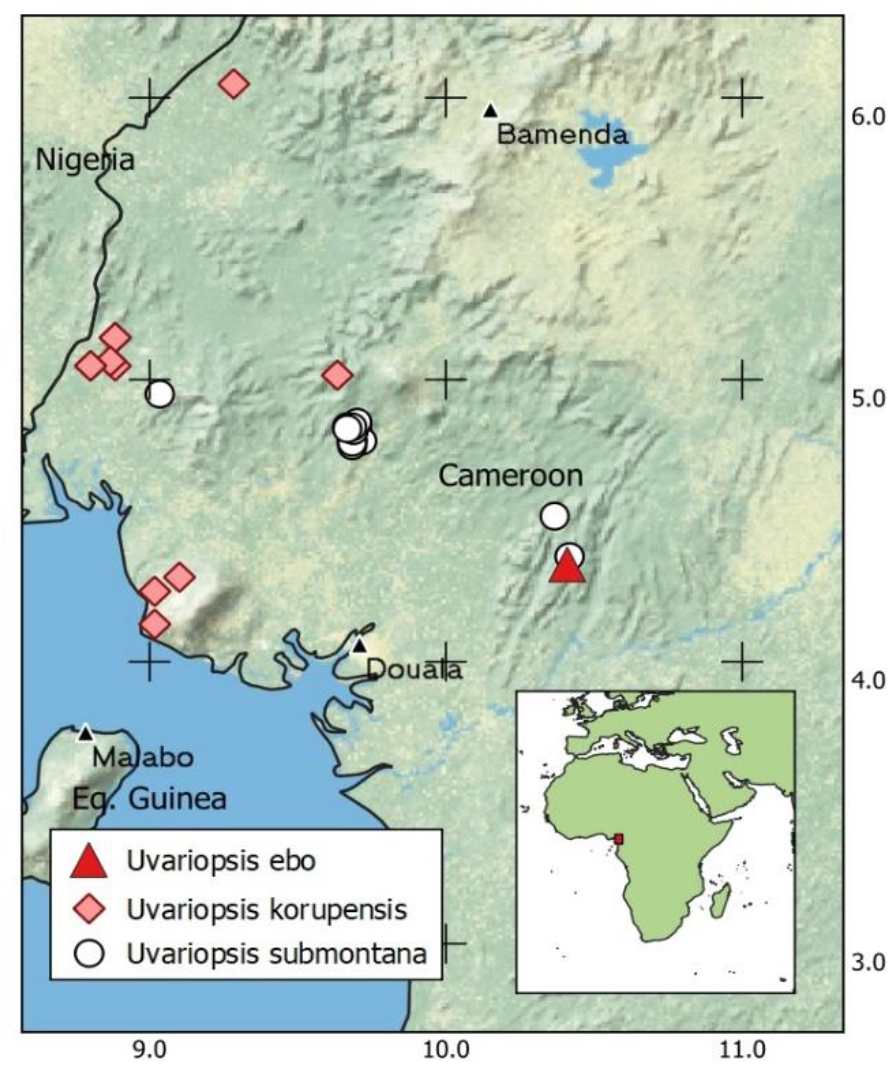
Global distribution of *Uvariopsis ebo*, together with *U. korupensis* and *U.submontana*.

The area of occupation of *Uvariopsis ebo* is estimated as 4 km^2^ using the IUCN preferred cell-size. The extent of occurrence is the same. In February 2020 it was discovered that moves were in place to convert the forest into two logging concessions (e.g. https://www.globalwildlife.org/blog/ebo-forest-a-stronghold-for-cameroons-wildlife/ and https://blog.resourceshark.com/cameroon-approves-logging-concession-that-will-destroy-ebo-forest-gorilla-habitat/ both accessed 19 Sept. 2020). Such logging would result in timber extraction that would open up the canopy and remove the intact habitat in which *Uvariopsis ebo* is found. Additionally, slash and burn agriculture often follows logging trails and would negatively impact the population of this species. Fortunately the logging concession was suspended in August 2020 due to representations to the President of Cameroon on the global importance of the biodiversity of Ebo (https://www.businesswire.com/news/home/20200817005135/en/Relief-in-the-Forest-Cameroonian-Government-Backtracks-on-the-Ebo-Forest accessed 19 Sept. 2020). However, the forest habitat of this species remains unprotected and threats of logging and conversion of the habitat to plantations remain, and mining is also a threat. *Uvariopsis ebo* is therefore here assessed as Critically Endangered, CR B1+2ab(iii), D.

#### PHENOLOGY

Flowering has only been observed in late March and early April (Bethan Morgan pers. comm. 2021).

#### ETYMOLOGY

Named as a noun in apposition for the forest of Ebo.

#### VERNACULAR NAMES & USES

None are known.

#### NOTES

An unusual and distinctive feature of *Uvariopsis ebo* is the shape and form of the corolla. The centre of the proximal half of each petal is concave before anthesis, forming a globose chamber for the staminal dome with the other three petals (Fig. 1). On the inner surface of the petal, the concave area is demarcated by an inverted U-shaped, distinct wall-like, broad ridge which seems to be the point of contact with those of the other sepals, sealing the chamber. The distal half of the petals and the margins of the proximal half are flat, wing-like and held against each other (applanate). In section therefore, the distal part of the corolla will appear cross-shaped (see Fig. 1). This seems to be an extreme form of the petal structure seen in the probably closely related Cameroonian species *Uvariopsis korupenis* and *U. submontana* which also occur in the SW quadrant of Cameroon (see Map 1). Additional species of *Uvariopsis* present at Ebo Forest are *U. zenkeri* Engl. and *U. submontana*.

## Discussion

### The range of *Uvariopsis ebo* and other endemic species in the Ebo Forest area

Abwe & Morgan, (2008) and Cheek *et al*. (2018a) give overviews of habitats, species and importance for conservation of the Ebo forest to which *Uvariopsis ebo* is restricted on current evidence. Sixty-nine globally threatened plant species are currently listed from Ebo on the IUCN Red List website and the number is set to rise rapidly as more of Cameroon’s rare species are assessed for their conservation status as part of the Cameroon TIPAs programme. The discovery of a new species to science at the Ebo forest is not unusual. Numerous new plant species have been published from Ebo in recent years. Examples of other species that, like *Uvariopsis ebo*, appear to be strictly endemic to the Ebo area on current evidence are: *Ardisia ebo* Cheek (Cheek & Xanthos, 2012), *Crateranthus cameroonensis* Cheek & Prance (Prance & Jongkind, 2015), *Gilbertiodendron ebo* Burgt & Mackinder, *Inversodicraea ebo* Cheek (Cheek *et al*. 2017), *Kupeantha ebo* M.Alvarez & Cheek (Cheek *et al*. 2018b), *Kupeanthaya bassi* M.Alvarez & Cheek (Alvarez *et al*. 2021), *Palisota ebo* Cheek (Cheek *et al*. 2018a) and *Pseudohydrosme ebo* Cheek (Cheek *et al*. 2021).

Further species described from Ebo have also been found further west, in the Cameroon Highlands, particularly at Mt Kupe and the Bakossi Mts (Cheek *et al*. 2004). Examples are *Myrianthus fosi* Cheek (in Harvey *et al*. 2010), *Salacia nigra* Cheek (Gosline & Cheek, 2014) and *Talbotiella ebo* Mackinder & Wieringa (Mackinder *et al*. 2010).

Additionally, several species initially thought endemic to Mt Kupe and the Bakossi Mts and adjoining areas in the Cameroon Highlands have subsequently been found at Ebo, e.g. *Coffea montekupensis* Stoff. (Stoffelen *et al*. 1997), *Costus kupensis* Maas & H. Maas (*Maas-van der Kamer et al. 2016), Deinbollia oreophila* Cheek (Cheek & Etuge 2009), *Microcos magnifica* Cheek (Cheek, 2017), and *Uvariopsis submontana* Kenfack, Gosline & Gereau (Kenfack *et al*. 2003). It is considered likely that additional Kupe species may yet be found at Ebo such as *Brachystephanus kupeensis* I.Darbysh. (Champluvier & Darbyshire, 2009), *Impatiens frithii* Cheek (Cheek & Csiba 2002) since new discoveries are still frequently being made in the Ebo Forest. Therefore, it is possible that *Uvariopsis ebo* might yet also be found in the Cameroon highlands, e.g. at Mt Kupe. However, this is thought to be only a relatively small possibility given the high level of survey effort at Mt Kupe: if it occurred there it is highly likely that it would have been recorded already since it is so spectacular it would be difficult to overlook.

## Conclusions

Such discoveries as this new species underline the urgency for making further such discoveries while it is still possible since in all but one of the cases given above, the species have very narrow geographic ranges and/or very few individuals, and face threats to their natural habitat, putting these species at high risk of extinction.

About 2000 new species of vascular plant have been discovered each year for the last decade or more. Until species are known to science, they cannot be assessed for their conservation status and the possibility of protecting them is reduced (Cheek *et al*. 2020). Documented extinctions of plant species are increasing, e.g. *Oxygyne triandra* Schltr. and *Afrothismia pachyantha* Schltr. of South West Region, Cameroon are now known to be globally extinct (Cheek & Williams 1999, Cheek *et al*. 2018c, Cheek *et al*. 2019). In some cases, species appear to be extinct even before they are known to science, such as *Vepris bali* Cheek, also from the Cross-Sanaga interval in Cameroon (Cheek *et al*. 2018d) and elsewhere, *Nepenthes maximoides* Cheek (King & Cheek, 2020). Most of the 815 Cameroonian species in the Red Data Book for the plants of Cameroon are threatened with extinction due to habitat clearance or degradation, especially of forest for small-holder and plantation agriculture following logging (Onana & Cheek, 2011). Efforts are now being made to delimit the highest priority areas in Cameroon for plant conservation as Tropical Important Plant Areas (TIPAs) using the revised IPA criteria set out in Darbyshire *et al*. (2017). This is intended to help avoid the global extinction of additional endemic species such as *Uvariopsis ebo* which will be included in the proposed Ebo Forest IPA.

With only one locality known, *Uvariopsis ebo* represents another narrowly endemic Cameroonian species threatened with extinction due to deforestation for oil palm plantations, small-scale agriculture, mining and logging (Onana & Cheek 2011, Cheek *et al*. 2018a).

## Acknowledgements

This paper was completed as part of the Cameroon TIPAs (Tropical Important Plant Areas) project at RBG, Kew, which is supported by Players of People’s Postcode Lottery. Ekwoge Abwe and Bethan Morgan and their team at the Ebo Forest programme are thanked hugely for making available the specimen and photos on which this paper is based and for expediting our botanical surveys in the Ebo forest of Cameroon over several years which allowed us to give context about the Ebo forest in this paper. We thank Janis Shillito for typing the manuscript, Lorna MacKinnon for the photos and specimen.

The heads of IRAD (Institute of Research in Agronomic Development)-National Herbarium of Cameroon, Yaoundé, successively Florence Ngo Ngwe, Eric Nana and Jean Betti Lagarde, are thanked for co-ordinating the co-operation with the Royal Botanic Gardens, Kew. The authors would like to thank two anonymous reviewers for comments on an earlier version of this manuscript.

